# Population level genetic memory of prior metabolic adaptation in *E. coli*

**DOI:** 10.1101/2022.09.20.508651

**Authors:** Sophia Katz, Claudia Grajeda-Iglesias, Bella Agranovich, Alia Ghrayeb, Ifat Abramovich, Sabrin Hilau, Eyal Gottlieb, Ruth Hershberg

## Abstract

Bacteria must often survive following the exhaustion of their external growth resources. Fitting with this need, many bacterial species that cannot sporulate, can enter a state known as long term stationary phase (LTSP) in which they can persist for years within spent media. Several recent studies have revealed the dynamics of genetic adaptation of *Escherichia coli* under LTSP. Yet, the metabolic consequences of such genetic adaptation were not addressed. Here, we characterized the metabolic changes LTSP populations experience and link them to their genetic adaptation. We observed that during growth within fresh resources *E. coli* produces the short chain fatty acid butyrate, which wildtype *E. coli* cannot consume. Once resources are otherwise exhausted, *E. coli* adapts genetically to consume butyrate through the convergent, temporally precise emergence of mutation combinations within genes that regulate fatty acid metabolism. These mutations appear to negatively affect bacterial fitness when butyrate is not available, and hence rapidly decrease in frequency, once all butyrate is consumed. Yet despite this, *E. coli* populations show a remarkable capability of maintaining a population-level genetic ‘memory’ of prior adaptation to consume butyrate. The maintenance of such a ‘memory’ allows bacteria to rapidly re-adapt, at an ecological, rather than an evolutionary timeframe, to re-consume previously encountered metabolites.

## Introduction

The ability of bacteria to survive within nutrient limited environments has fascinated the scientific community for the past 90 years (Zobell and Grant 1943). This ability results from bacterial evolution in natural environments, where bacteria are likely repeatedly exposed to short periods of feast and long periods of famine. Bacteria pursue several strategies to face long periods of starvation. Some bacterial species can undergo sporulation, allowing them to maintain complete or near complete dormancy for many years. While sporulation has rightfully received much attention, most known bacteria species cannot sporulate, yet many can still survive for years following the exhaustion of their nutrients, by entering a state termed long term stationary phase (LTSP) (Zambrano, et al. 1993; Finkel 2006). In LTSP a sub-population of cells can survive by recycling the remains of their deceased brethren.

We and others have previously reported on evolutionary experiments designed to probe the dynamics of *E. coli* genetic adaptation under LTSP (Avrani, et al. 2017; Chib, et al. 2017; Avrani, et al. 2020; Gross, et al. 2020; Katz, et al. 2021; Ratib, et al. 2021). In one such experiment, initiated in July 2015 and ongoing until this day, we established five independent populations by inoculating *E. coli* cells into rich media and incubating them within a shaking incubator. ∼10 clones from each of these five populations were sequenced, at nine time points, spanning the first three years of the experiments. Results of these experiments thus far show that *E. coli* LTSP populations rapidly adapt genetically, in an extremely convergent manner, through mutations to several important regulatory and metabolic genes (Avrani, et al. 2017; Katz, et al. 2021).

The first adaptation we observed to occur in *E. coli* under LTSP, involves mutations within highly specific sites of the RNA polymerase core enzyme (RNAPC) (Avrani, et al. 2017). A second, highly convergent adaptation that emerges with great temporal precision involves mutations to fatty acid metabolism genes. Specifically, at day 22 of our experiments we observed that, across independently evolving populations, ∼80% of sequenced clones carry, in addition to the RNAPC adaptation, a mutation in the *fadR* gene which encodes a master regulator of fatty acid metabolism (van Aalten, et al. 2000), combined with a second mutation in either *atoC* or *atoS*, that together encode a two-component system that regulates fatty acid metabolism (Lioliou and Kyriakidis 2004). The adaptations within *fadR* and *atoC*/*atoS* that appear with high convergence at day 22 were not seen at the previous (day 11) or subsequent (day 32) sampled time points (Avrani, et al. 2017).

The ability of bacteria to enter LTSP and persist in the absence of external nutrients for many years constitutes a metabolic mystery. Yet the metabolic particulars of LTSP have not been addressed. Here we provide the first characterization of the metabolites present within LTSP exhausted media, over the first 32 days under LTSP. This in turn allowed us to ascertain the function of the *fadR* + *atoC*/*atoS* adaptations, demonstrating that these enable the consumption of the short chain fatty acid butyrate, which we show *E. coli* produces during the first 24 hours of growth in rich media. We could further show that despite these adaptations demonstrating an apparent cost, and despite our populations being well mixed, *E. coli* LTSP populations are able to maintain minorities of cells carrying these adaptations, even when no butyrate is present. Such maintenance allows populations that adapted to consume butyrate, to re-adapt to consuming this metabolite much more rapidly than populations that never encountered butyrate before.

## Results

### Characterization of the metabolites present within LTSP media as a function of time spent under LTSP

To examine the metabolic consequences of LTSP adaptations, we initiated new LTSP experiments and measured changes of different metabolites in the media over 32 days (Materials and Methods). Sugars serving as main carbon sources such as glucose and its disaccharide trehalose (a major sugar component of LB (Puentes-Tellez, et al. 2013)) were rapidly consumed, following three and 24 hours of growth respectively (**Figure 1A and Supplementary Table S1**). In contrast, uptake of fructose, and of sugar alcohols, considered to be metabolic products (Hanko and Rohrer 2000), was minimal even following 32 days. A significant increase in the concentration of the pentoses, ribose and xylose was observed towards day 32, likely due to the breakdown of dying cells and to the deficient utilization of these sugars, relative to other available carbon sources (Ammar, et al. 2018; Fujiwara, et al. 2020). Amino acids, that are relatively not bioenergetically costly to produce, such as glycine, serine, and threonine, were rapidly consumed. In contrast, very little, if any uptake occurred for phenylalanine, histidine, and the branched-chain amino acids leucine, isoleucine and valine (**Figure 1B and Supplementary Table S2**), all of which are much more costly to produce (Kaleta, et al. 2013). This observation fits well with previous data suggesting a compromise between the cost and the potential energetic benefit of degrading amino acids (Zampieri, et al. 2019).

**Figure 1.**
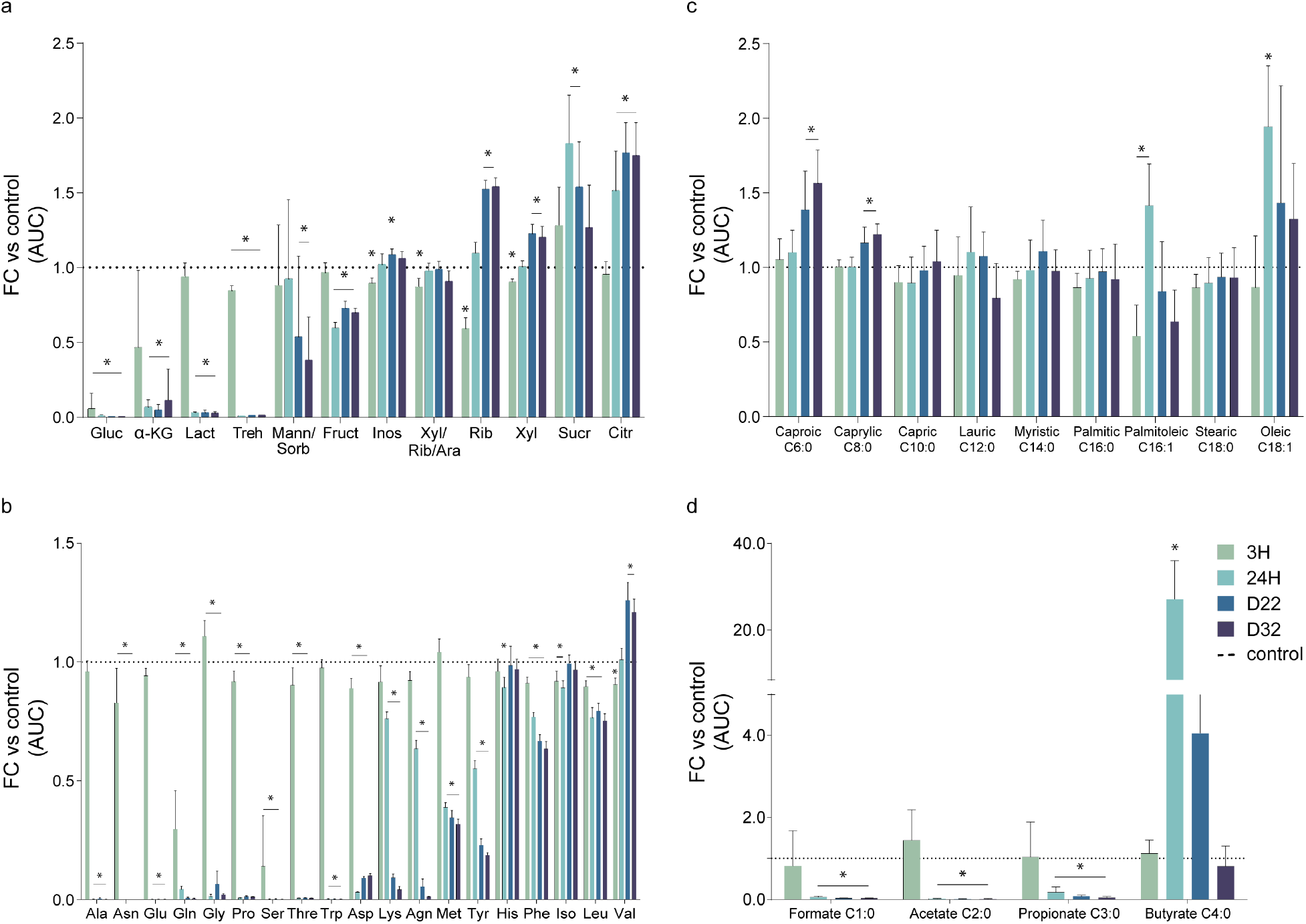
Changes in the presence of metabolites within LTSP media, during the first 32 days of adaptation. Data obtained from five to nine independent LTSP populations, which were established as described in the text. For each population, consumed and excreted metabolites were identified in the media, following 3 h, 24 h, 22 days and 32 days of culture. Six flasks containing only LB without bacteria were used as controls. The fold change (FC) area under the curve (AUC), compared to the controls, was calculated for each identified metabolite at the indicated time point. (A) Glucose and other sugars and derivatives, (B) proteogenic amino acids, and (C) medium- and long-chain free fatty acids were analyzed by targeted LC-MS. (D) Short-chain fatty acids were identified by targeted GC-MS, previous derivatization. Mean values with error bars representing standard deviations around the mean are shown. Statistical significance between two samples was calculated using a Mann-Whitney two-sided test, to compare the absolute values from each timepoint versus control. * = P <0.05. (The detailed data and their statistical analyses can be found in **Supplementary tables S1-S3**).

### E. coli populations produce butyrate during initial growth in fresh rich media and adapt to consume it through the acquisition of mutations within *fadR* and *atoC*/*atoS*

The concentration of medium- and long-chain free fatty acids did not significantly reduce with time (**Figure 1C and Supplementary Table S3**). However, the short chain fatty acids (SCFA), formate (C1), acetate (C2) and propionate (C3), were consumed by 24 hours (**Figure 1D and Supplementary Table S3**). Intriguingly, The SCFA butyrate (C4) was massively produced during the first 24 hours of growth and was then consumed in a temporally precise manner following 18-22 days of culture (**Figure 2A**). Butyrate serves as the main energy source for colonocytes (Bultman 2016). Yet, despite its importance to homeostasis, the human body cannot synthesize butyrate and relies on anaerobic fermentation by commensal gastrointestinal bacteria, of the phyla Bacteroidetes and Firmicutes (Zhang, et al. 2009). Wildtype *E. coli* is not known to produce butyrate ()Jawed, et al. 2016(or to consume it (Overath and Raufuss 1967). Accordingly, the ancestral *E. coli* K12 MG1655 strain, used to initiate our experiments, is not able to consume butyrate (**Figure 2B**). However, our experiments with LTSP evolved clones, carrying the *fadR* + *atoC*/*atoS* mutations demonstrated their acquired ability to consume butyrate (**Figure 2B**). Population samples from day 22, in which these butyrate consuming clones are present at high frequencies can also consume butyrate, while samples extracted from day 11 and day 32 of LTSP cannot (**Figure 2B**). Taken together, these results suggest that LTSP populations rapidly adapt to consume the butyrate they generated during the first day of growth and that they do so through the convergent acquisition of a specific combination of mutations. Once butyrate is fully consumed, the *fadR* + *atoC*/*atoS* mutations are no longer useful and they undergo sharp reductions in frequencies by day 32 of our original experiment (Avrani, et al. 2017). That this reduction in frequency occurred so rapidly, across multiple populations, suggests that in the absence of butyrate, these mutations come at a relative cost.

**Figure 2.**
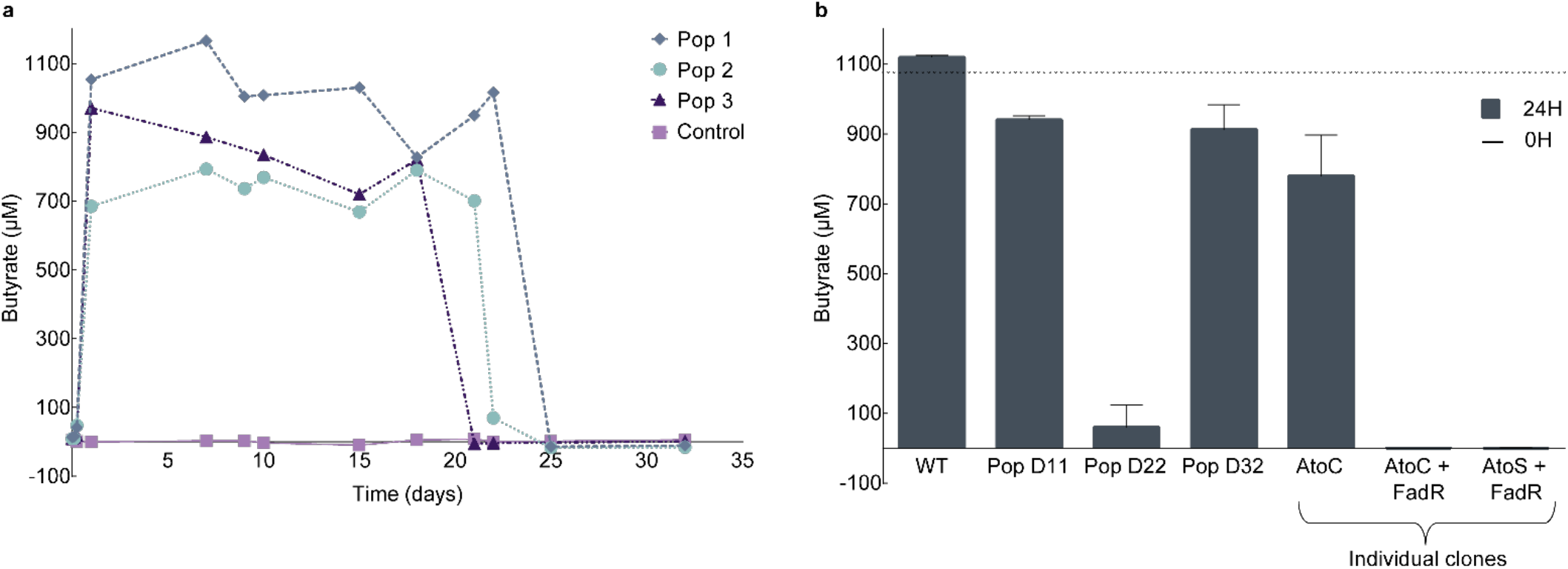
Butyrate kinetics. **(A) Butyrate is produced during the first 24 hours of growth in LB and then consumed at days 18-22**. Butyrate kinetics were monitored for 32 days by GCMS analyses on three independent LTSP populations, with LB media without inoculated bacteria serving as a control (purple). **(B) day 22 LTSP population samples and clones carrying *fadR* + *atoC* or *fadR* + *atoS* mutation combinations can consume butyrate**. The ability to consume butyrate was examined for each sample or clone, through GCMS measurements of the levels of butyrate remaining after 24 hours. Mean values across three biological replicas, per sample or clone, are presented with error bars representing standard deviations around the mean. The levels of butyrate added at day 0 are marked with a dotted line.

### Butyrate consumption adaptations provide a growth advantage when butyrate is provided, but are rapidly outcompeted once butyrate is no longer available

Next, we sought to examine what would happen if butyrate was continuously available. We established six new LTSP populations. For three of them we supplemented the media daily, from day 18 to day 55, with 1 mM butyrate (a concentration similar to that produced during the first 24 hours of growth (**Figure 2A**). The remaining three populations served as controls and were allowed to evolve without further intervention. In two of the three butyrate-supplemented populations, the butyrate produced during the first 24 hours of growth was consumed at days 17 and 18 of the experiment, and the supplemented butyrate was then continuously consumed daily (**Figure 3A**). Starting at day 18 and up to day 55, bacteria from these two butyrate-supplemented populations were able to grow substantially more than in the absence of butyrate, reflected in much higher colony forming unit (CFU) counts (**Figure 3B**). In the third population, butyrate produced during the first 24 hours of growth was consumed earlier, at day seven, and butyrate supplemented Starting at day 18, initially accumulated, and was only consumed from day 26 onwards (**Figure S1A**). In line with this, the increase in CFU was observed for this population only from day 26 (**Figure S1B**). Combined, these results demonstrate that LTSP bacteria that adapt to consume butyrate can grow on butyrate when it continuously made available.

**Figure 3.**
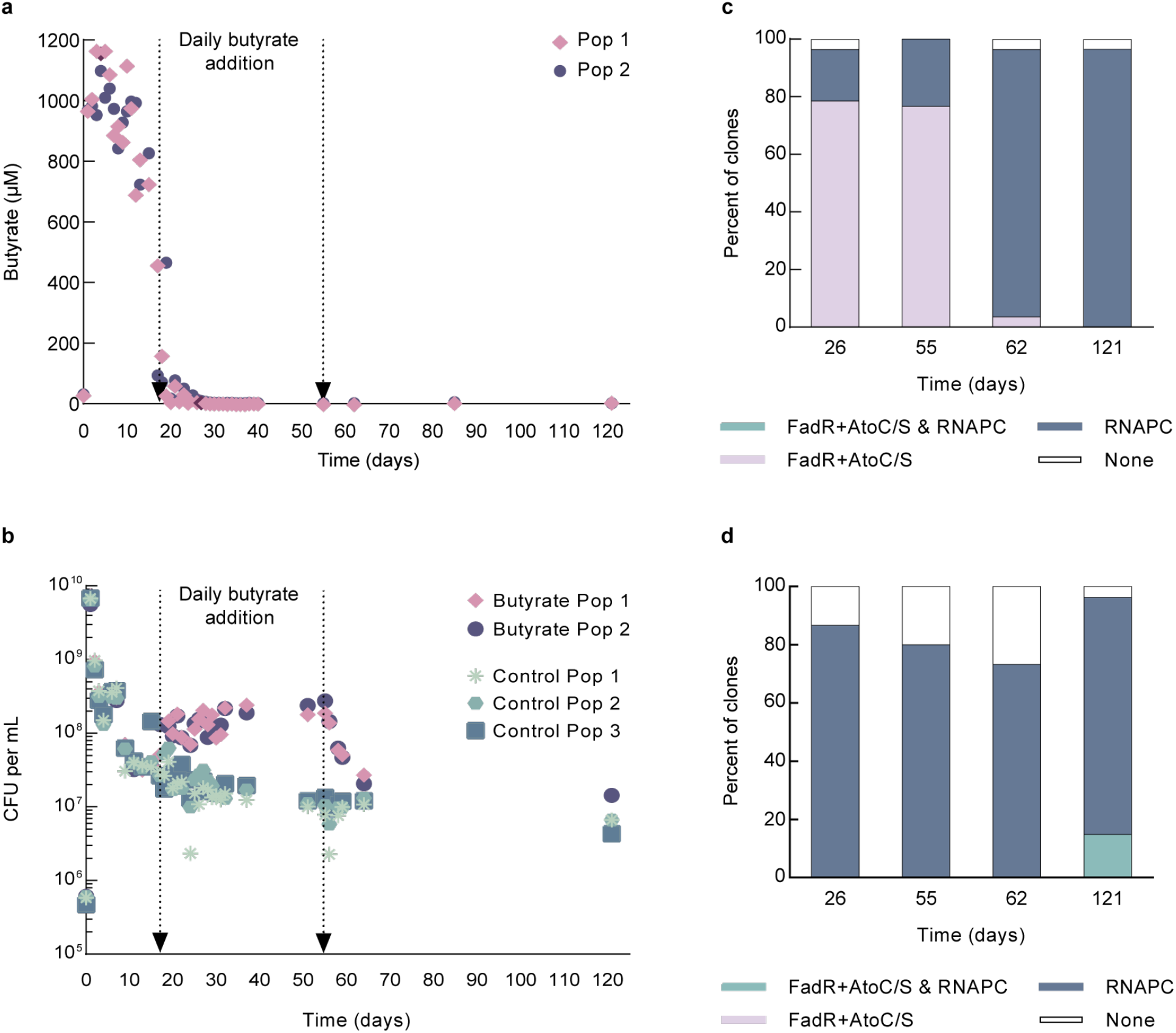
Daily supplementation of butyrate and its effect on LTSP populations. Six LTSP populations were established. To three of the six populations, butyrate was added daily starting at day 18 and ending at day 55. **(A)** Levels of butyrate measured within two populations to which butyrate was added, as a function of time. The time range in which butyrate was added is marked on the graph (A similar plot for the third population is presented in **Supplementary Figure S1A**). **(B)** Mean number of viable cells with time in the two LTSP populations supplemented daily with butyrate (pink/purple) or in control LTSP populations which were allowed to evolve without interference (green). Numbers of viable cells were estimated by counting colony-forming units (CFU). The time range in which butyrate was added is marked on the graph. (A similar plot for the third population is presented in **Supplementary Figure S1B**). **(C)** In the three populations to which butyrate is added between days 18 and 55, a high frequency of clones sequenced at day 26 and 55 carry butyrate consumption adaptations but do not carry any adaptations within the RNAPC. Clones sequenced at day 62 very rarely carry butyrate consumption adaptations and often carry an RNAPC adaptation. **(D)** Control populations to which butyrate was not added almost never contain butyrate consumption adaptations at observable frequencies. In contrast, these populations contain high frequencies of clones carrying RNAPC adaptations at all time points.

We fully sequenced 9-10 clones from each of the six populations at each of four time points: days 26, 55, 62 and 121. At days 26 and 55, clones carrying the *fadR* + *atoC*/*atoS* mutation combination were seen at high frequencies (ranging from 66.7% to 100%) within all three populations supplemented with butyrate (**Figure 3C**, and **Figure S1C Supplementary table S4**). Strikingly, in each of the three populations, more than a single genotype carrying these mutation combinations appeared to segregate simultaneously (**Figure S1C**), highlighting the remarkably extensive genetic variation present within LTSP populations. In contrast, no butyrate consumption genotypes were observed at days 26 or 55 within the control populations (**Figure 3D**, and **Figure S1D Supplementary table S4**). In our original experiments, the *fadR* + *atoC*/*atoS* mutation combination was found within day 22 clones that also carried an RNAPC adaptation (Avrani, et al. 2017). However, all clones sequenced from our butyrate-supplemented populations carried either the *fadR* + *atoC*/*atoS* mutations, or an RNAPC adaptation or, in rarer cases, neither of the two (**Figure 3C** and **Supplementary table S4**). We have previously shown that LTSP RNAPC adaptations severely reduce growth rates in rich media, where nutritional resources are readily available (Avrani, et al. 2017; Avrani, et al. 2020). This may explain why these RNAPC adaptations would be disfavored, in butyrate supplemented LTSP populations, in clones that have adapted to be able to grow on butyrate. Yet, clones that carry an RNAPC adaptation, but do not carry mutations enabling them to consume butyrate still persist within our populations at sufficiently high frequencies to allow their detection through the sequencing of a handful of clones out of ∼400*10^7^ cells present within our populations. This suggests that these clones either occupy a niche that provides them with an advantage despite their inability to consume butyrate, or that they are maintained at high frequencies for more than five weeks, despite carrying substantial costs to fitness. In contrast, once butyrate supplementation stops (at day 55), clones carrying the butyrate consumption adaptations reduce in frequencies rapidly, so by day 62, no such clones were observed in two of the butyrate-supplemented populations and only a single such clone was detected in the third population (**Figure 3C** and **Figure S1C Supplementary table S4**). This suggests that butyrate consumption adaptations are associated with costs that rapidly drive their frequencies down once butyrate is no longer available.

### Despite apparent costs, LTSP populations are able to maintain a relatively long standing ‘memory’ of their prior ability to consume butyrate

Bacteria often exist within fluctuating environments, in which a metabolite that became available once, is more likely to become available again later. Being able to rapidly re-adapt to consume a metabolite that a population already adapted to consume once before, may thus be beneficial. Yet, as seems to be the case for the butyrate-consumption adaptations, adapting to consume a new metabolite may often come at a cost to fitness. We wanted to understand whether despite the apparent costs associated with the butyrate-consumption adaptations, populations are able to maintain a genetic population-level ‘memory’ of their previous adaptations to consume butyrate. In other words, we aimed to examine whether populations that once grew in the presence of butyrate and adapted to consume it, would re-adapt more rapidly to consume it, once it was again available. To test this, we needed to obtain control populations that were never exposed to butyrate. Towards this end, we established six new LTSP populations, as usual starting by growing cells in LB. After 24 hours of growth, we filtered the bacteria out of their media, washed and re-inoculated them into M9 minimal media (that contains no butyrate and no other carbon sources). To three of the six populations we added butyrate at similar concentrations to those produced by *E. coli* during the first 24 hours of growth in LB (1 mM). Butyrate levels were monitored for all six populations. As expected, no butyrate was detected for the control populations. The M9 populations to which butyrate was added, consumed it earlier than LB-maintained populations, at day six for one of the M9 populations and day 11 for the remaining two (**Figure 4A**). Once butyrate was consumed from these three populations, we waited an additional four weeks and then sampled each of the six populations and added butyrate to each of the samples, either once (**Figure 4**), or daily for seven days (**Figure S2**). As shown in **Figure 4B and Figure S2B**, the populations that never saw butyrate, could not consume it for the seven days examined. In contrast, the populations that had seen butyrate and had adapted to consume it, re-adapted to consume it rapidly, doing so within two to five days (**Figure 4A** and **Figure S2A**).

**Figure 4.**
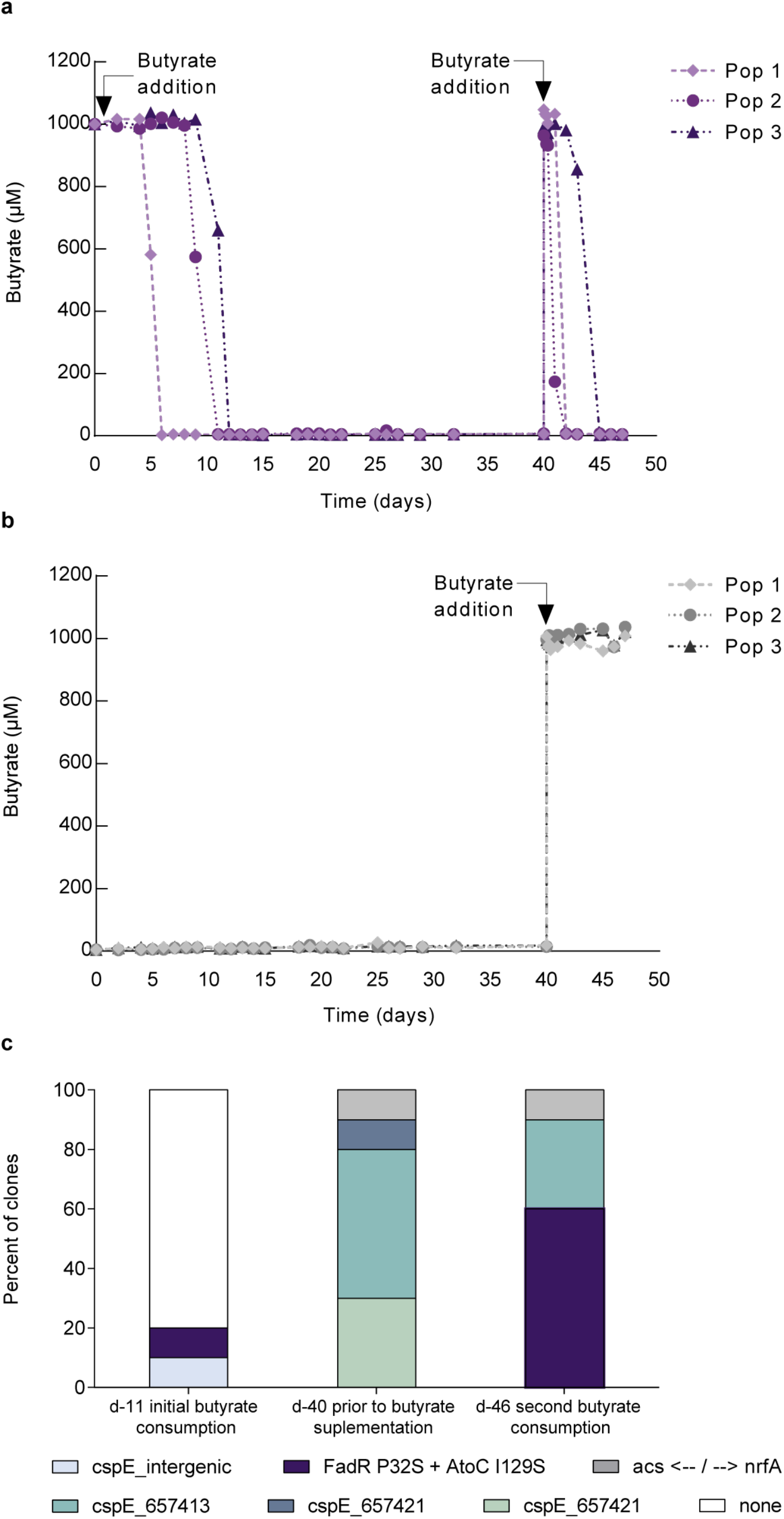
Populations that previously encountered butyrate adapt more rapidly to consume it, once it again becomes available. One day after initiation, cells from three LTSP experiments were filtered out of their LB media and re-inoculated into three flasks of M9 minimal media, either supplemented with 1 mM butyrate **(A)**, or not **(B)**. When butyrate was provided, it was initially consumed by day 12. Following four additional weeks, populations were sampled and assayed for their ability to consume butyrate. As shown, only populations that initially received butyrate could then consume it, during the week following its addition into their media (**A**, arrows represent time points in which butyrate was added). **(C)** Sequencing data from butyrate-supplemented population 3 reveals that the same mutation combination was responsible for initial (day 11) and secondary (day 46) butyrate consumption. Yet, this mutation combination was not present at observable frequencies at day 40, immediately prior to the second supplementation with butyrate. The full list of mutations identified can be found in **Table S5**.

Focusing on the populations that initially received butyrate and then received it again only once, we sequenced 9-10 clones from each population at the time point in which they consumed the butyrate for the second time. In two of the three populations 33.3% to 60% of the clones sequenced carried a *fadR* + *atoC*/*atoS* mutation combination (**Supplementary table S5**). Furthermore, in one of these two populations we could also see the exact same mutation combination at the time at which that population initially consumed the butyrate (day 11 of the experiment (**Figure 4C Supplementary table S5**)). Clones carrying the butyrate-consumption genotype were not observed when sequencing 10 clones from the sampling time-point, prior to the addition of butyrate (**Figure 4C Supplementary table S5**). Combined, this demonstrates that the genotypes that arise in frequency to consume the butyrate initially, decrease in frequency, so they can no longer be observed four weeks later, but are still maintained within their populations at lower frequency and can thus again increase in frequency to consume the butyrate the second time. No mutations of note were identified when sequencing ∼10 clones from the other populations at the time they initially consumed the butyrate (**Supplementary table S5**). For the third population we did not observe any clones carrying the *fadR* + *atoC*/*atoS* mutation combination, even once they consumed the butyrate for the second time.

Our inability to always identify clones carrying the *fadR* + *atoC*/*atoS* mutation combination when butyrate is being consumed, may be explained by the fact that we sequenced only 9-10 clones out of a population of ∼10^8^. This means that any mutation that is segregating at slightly lower frequencies might not be detectable. Nevertheless, combined our results here demonstrate that despite the costs associated with the butyrate-consumption adaptations, they remain within our populations for at least four weeks, following the consumption of butyrate. This in turn allows the populations that previously adapted to consume butyrate to establish a population-level genetic ‘memory’ of their prior adaptation, and more rapidly re-adapt to consume butyrate, once it again becomes available.

## Discussion

Wildtype *E. coli* cannot consume the short chain fatty acid butyrate. Here, we show that *E. coli* produces butyrate during the first 24 hours of growth in fresh LB. This butyrate remains within the media until, in a remarkably convergent and temporally precise manner, LTSP clones adapt to consume it, through the acquisition of mutation combinations within the *fadR* and *atoC*/*atoS* genes. These adaptations allow *E. coli* to grow on butyrate, when it is daily supplemented into the LTSP media. However, once supplementation stops and butyrate is no longer available, the frequency of the butyrate consumption genotypes convergently reduces within a very short period of time, strongly suggesting that these adaptations carry a cost. Despite this apparent cost, and despite the fact that our populations are well mixed, and that previous studies have demonstrated them to be subject to very strong selection (Avrani, et al. 2017; Katz, et al. 2021), butyrate consumption genotypes remain at lower frequencies within our populations for a substantial period of time. This in turn, allows these populations to re-adapt more rapidly to consume butyrate, once it again becomes available, compared to populations that never encountered butyrate before, and never adapted to consume it.

It has long been understood that environmental heterogeneity can support the long-term maintenance of different genotypes, each of which is specialized to a different set of conditions (reviewed in (Hedrick, et al. 1976; Kassen 2104)). At the same time, within well mixed populations, of the type studied here, for which conditions change only temporally, the co-existence of specialist genotypes is thought to be more limited, and possible only under specific sets of circumstances (Hedrick 1976; Reboud and Bell 1997; Kassen 2002, 2104). Our results indicate that even for such well-mixed populations genotypes specialized towards previously encountered conditions may be maintained for substantial periods of time. It is possible that our experimental design that, similar to most natural environments, does not dilute out slow growing or dormant cells, enables this maintenance. While most evolutionary experiments rely on continuous culturing or on serial dilution (reviewed in (Barrick and Lenski 2013; Gresham and Dunham 2014; Kassen 2104)), which do not allow variation in growth rates to freely persist within the studied populations, our experiment does not artificially limit such variation (Katz, et al. 2021). Dormant spores or bacterial cells that enter a dormant or extremely slow growing state are at least partially immune from selection (Kussell, et al. 2005; Jones and Lennon 2010; Gray, et al. 2019), allowing them to survive despite carrying deleterious genotypes. Dormancy or extreme slow growth is thus one manner in which populations may be able to maintain within them non-favored genotypes, specialized towards a previously encountered condition (Miller and Klausmeier 2017; Shoemaker and Lennon 2018). Balancing selection can also enhance the co-existence of specialist genotypes (Good, et al. 2017; Turner, et al. 2020). Enabling populations to maintain higher variation in growth rate may increase the opportunities for such balancing selection (Levy, et al. 2012; Merot, et al. 2020) and may therefore enhance the ability of populations to maintain within them longer-term genetic variation, including genotypes adapted to previous conditions.

Importantly, our study revealed that the previous metabolic conditions toward which the bacteria are adapted to under LTSP were, in fact, generated by the bacteria themselves. Butyrate is produced and secreted to the media at times of nutritional abundance, and it forms a metabolic pressure selecting for genetic adaptations that enable later butyrate consumption under LTSP. It is therefore plausible that once bacteria are adapted to thrive on a nutritional resource that was initially produced by the same population, they are likely to maintain genetic memory for such conditions, even if it comes with a cost once the nutrient is fully consumed.

That large *E. coli* LTSP populations can maintain a population-level genetic ‘memory’ of their previously acquired ability to consume a particular metabolite, may very well reflect a broader ability of many bacterial populations to “remember” their previous metabolic adaptations. When such a ‘memory’ is maintained, genetic adaptation to fluctuating conditions can occur at an ecological rather than an evolutionary time scale. Establishment of a genetic population-level ‘memory’ of adaptations to consume metabolites that are periodically encountered within a certain environment could also enable bacterial populations to obtain a better ‘foot-hold’ within their environment, by offering a type of protection from invasion by outside populations. After all, populations that already established a ‘memory’ of the metabolites which are most commonly available within a particular environment, will have an advantage in their ability to re-adapt more rapidly to consume these metabolites, compared to an entirely naïve external population.

## Materials and Methods

### Media preparation

Luria-Bertani (LB) medium was prepared with 10 g L^−1^ tryptone (BD), 5 g L^−1^ yeast extract (BD), and 5 g L^−1^ NaCl. For (LB) agar plates, 15 g L^−1^ agar (BD) were added to the same formulation. Minimal medium (M9) was prepared according to manufacturer instructions (Difco™ M9 Minimal Salts, 5× Cat. No. 248510), and supplemented with 2 mL L^−1^ MgSO_4_ 1 M, and 0.1 mL L^−1^ CaCl_2_ 1 M. Butyrate stock solution (Sigma Aldrich CAS no. 107-92-6) was prepared in ultra-pure water at a concentration of 100 mM, and added for a final concentration in the culture medium of 1 mM. All media and solutions were either filter sterilized (0.45 μm pore size) or autoclaved before use.

### LTSP experiments

Single colonies of *E. coli* K12 MG1655 (WT, ancestor) were grown in test tubes containing 4 mL of fresh LB medium in a shaking incubator (225 rpm at 37 °C), until they reached an OD ∼ 0.4. Then, 1 mL was taken to a 2 L-polycarbonate breathing flask containing 400 mL of fresh LB, for a final concentration of ∼ 2×10^6^ cells/mL. Flasks were placed in an incubator set at 37°C, with shaking at 225 rpm. No new nutrients or resources were added to the cultures with time, except for sterile water that was added to compensate for evaporation every 10–15 days, according to the weight lost by each flask during that time period.

Samples from the culture media were taken at regular intervals, centrifuged (5 min at 5000*g*/ 4°C), and the supernatant was stored at -20°C for further metabolomics studies. To estimate bacterial viability, 1 mL of each culture was sampled. Dilutions were plated using a robotic plater to evaluate viability through live counts. Samples were stored in 50% glycerol at -80°C for future analysis.

<u>For the daily supplementation of butyrate experiment,</u> single colonies were allowed to recover in 4 mL-fresh LB in a shaking incubator (225 rpm at 37 °C), until an OD ∼ 0.4 was reached. From each of three ancestral clones, two independent LTSP experiments were started (for a total of six experiments), as described, by inoculating *E. coli* K12 MG1655 into 400 mL-fresh LB medium, to a final concentration of ∼ 2×10^6^ cells/mL. Flasks were kept in a shaking incubator at 37°C 225 rpm. Starting at day 18 of these experiments, three populations (each one from a different ancestor) were supplemented daily with 1 mL of butyrate (for a final concentration of 1mM), up to day 55. For each butyrate-supplemented population, an additional flask with no supplementation was kept as a control. Population sampling was done regularly.

<u>For the M9 experiments,</u> single colonies were allowed to recover in 4 mL-fresh LB in a shaking incubator (225 rpm at 37 °C), until an OD ∼ 0.4 was reached. From each of three ancestral clones, 2 independent LTSP experiments were started (for a total of six experiments), as described, by inoculating *E. coli* K12 MG1655 into 400 mL-fresh LB medium, to a final concentration of ∼ 2×10^6^ cells/mL. Flasks were kept in a shaking incubator at 37°C/ 225 rpm during 24 h. Next, the total population in each flask was harvested by centrifugation (20 min at 4000g) and rinsed once in M9. All populations were resuspended in 400 mL of M9, three of them (each one from a different ancestor) received butyrate at a final concentration of 1 mM. At day 40 of the LTSP-M9 experiments, 20 mL of each of the six populations were withdrawn and split into two 50 mL-Erlenmeyer flasks (10 mL per flask). Each flask received either a single or a daily dose of butyrate (1 mM final concentration). The Erlenmeyer cultures were kept in a shaking incubator at 37°C/ 225 rpm, for one week. Culture media samples and CFU measurements were conducted as described above.

### Monitoring butyrate consumption

In order to monitor the consumption of butyrate, Frozen cultures from our previous study(Avrani, et al. 2017), of *E. coli* K12 MG1655 (WT, ancestor), LTSP populations 1,2 and 4 at days 11, 22 and 32, and isolated colonies from the clones summarized in **Supplementary table S6**, were thawed and allowed to grow overnight in liquid LB. Additionally, the wildtype ancestral strain was also thawed and grown overnight. At the following day, each type of resulting culture was diluted 1:50 into fresh LB media and recovered for an additional two hours, until they reached an OD ∼ 0.2. Cells were harvested by centrifugation (5 min at 5000*g*) and rinsed twice in M9 media. The cells were then resuspended in 5 mL of M9 (baseline) or M9 supplemented with butyrate (1mM final concentration). Tubes were placed in a shaking incubator (225 rpm at 37 °C) and sampled at time 0, 3 and 24 hours. Culture media samples were obtained after cell centrifugation (5 min at 5000*g*/ 4°C) and stored at -20°C for further metabolomics analysis.

### Profiling short-chain fatty acids (SCFA) by GCMS

Representative metabolite standards were used for quality control. ^13^C-labeled isotopes of formate, propionate, butyrate, and ^2^H_3_-acetate, were used as internal standards for quantification.

#### Chemical derivatization for SCFA

The chemical derivatization procedure of samples using ethyl chloroformate (ECF) was adapted from the protocol described in(Tumanov, et al. 2016). Briefly, 200 μL of medium sample was added to a 1.5 mL microfuge tube, followed by addition of 10 μL of internal standard, 10 μL of benzyl alcohol, 10 μL of sodium hydroxide 1 M, and 50 μL of pyridine. The tube was then placed on ice for 5 min. Next, 20 μL of ECF were added immediately followed by vigorous vortexing for 30 s. As gas builds up in the microfuge tube during the derivatization reaction, it is very important that samples are cold enough. Tubes may be carefully opened in the middle of the vortexing period to relieve pressure and put back to vortex to complete the required length of time. After vortexing, 200 μL of MTBE were added, the sample vortexed for another 20 s, and centrifuged at 10,000*g* or max speed for 5 min. 120 μL microliters of the resulting upper layer were transferred to a GC vial with insert for analysis.

#### SCFA detection and quantification

Derivatized samples were analyzed with an Agilent 7890B GC system coupled to a 7000 Triple Quadrupole GC-MS system. The column was Phenomenex ZB-1701 column (30 m x 0.25 mm x 0.25 μm), with the following oven program: initial temperature 50°C/ hold 2 min; increment at 10°C/min to 140°C/ hold 0 min; increment at 20°C/min to 182°C/ hold 1 min; and increment at 50°C/min to 280°C/ hold 0 min. Run time 16.1 min. Samples (1 μL) were injected using split mode (20:1, 28 mL/min split flow). The column gas flow was held at 1.4 mL/min of He. The temperature of the inlet was 280°C, the interface temperature 230°C, and the quadrupole temperature 150°C. The column was equilibrated for 2 min before each analysis. The mass spectrometer was operated in single-ion monitoring (SIM) mode between 9.9 and 14.0 min. A segment for each compound was defined: benzyl-formate at 9.9 min, fragments *m/z* 136 and 137; benzyl-acetate at 11.0 min, fragments *m/z* 150, 151, 152 and 153; benzyl-propionate at 12.2, fragments *m/z* 164, 165, 166 and 167; and benzyl-butyrate at 13 min, fragments *m/z* 178,179,180, and 182. All segments at 2.5 cycles/s. Agilent MassHunter Workstation Software (version B.07.01 SP1) was employed for automated data processing, using peak area for absolute concentrations calculations.

### Metabolites profile by LCMS

#### Metabolite extraction

For the extraction of secreted metabolites, 50 μL of medium was diluted in 950 μL of ice-cold extraction solvent (50% methanol, 30% acetonitrile, 20% water), vortexed at high speed/ 4°C during 10 min, centrifuged at 16 000*g* for 10 min at 4°C. Supernatant was transferred into LC-MS glass vials and stored at −80°C until measurement. For the extraction of intracellular metabolites cell pellets were vortexed with 200 μL of ice-cold extraction solvent during 10 min, and incubated at -20°C for another 10 min. These cold-vortex / incubating steps were repeated three times. Pellets were subsequently centrifuged at 16000*g* for 10 min at 4°C, and the supernatant were transferred into LC-MS glass vials with inserts and stored at −80°C until measurement.

#### LC-MS measurements

LC-MS metabolomics analysis was performed as described previously(Mackay, et al. 2015; Meiser, et al. 2016) (Mackay et al., 2015; Meiser et al. 2016). Briefly a Q Exactive Orbitrap mass spectrometer (Thermo Fisher Scientific) was used, together with a Thermo UltiMate 3000 high-performance liquid chromatography (HPLC) system. The HPLC setup consisted of a ZIC-pHILIC column (SeQuant, 150 mm x 2.1 mm; Merck KGaA, 5 mm), with a ZIC-pHILIC guard column (SeQuant, 20 mm x 2.1 mm) and an initial mobile phase of 20% of 20 mM ammonium carbonate (pH 9.2) and 80% acetonitrile. Cell and medium extracts (5 μL) were injected, and metabolites were separated over a 15-min mobile phase gradient, decreasing the acetonitrile content to 20%, at a flow rate of 200 μL/min and a column temperature of 45°C. The total analysis time was 27 min. All metabolites were detected across a mass range of 75 to 1 000 mass/charge ratio (*m/z*) using the Exactive mass spectrometer at a resolution of 35 000 (at 200 *m/z*), with electrospray ionization and polarity switching to enable both positive and negative ions to be determined in the same run. Lock masses were used, and the mass accuracy obtained for all metabolites was below 5 ppm. Data were acquired using Thermo Xcalibur software. The peak areas of different metabolites were determined using Thermo TraceFinder 4.1 software, where metabolites were identified by the exact mass of the singly charged ion and by known retention time on the HPLC column, using an in-house MS library built by running commercial standards of all metabolites detected.

### DNA extraction, sequencing library preparation and mutation calling

Frozen cultures were thawed and dilutions were plated and grown over night. Colonies to be sequenced were used to inoculate 4 ml of LB medium in a test tube and were grown until they reached an OD of 1. One milliliter of the culture was centrifuged at 5,000 g for 5 min and the pellet was used for DNA extraction. The remainder of each culture was then archived by freezing in 50% of glycerol at -80°C. DNA was extracted using the Macherey-Nagel NucleoSpin 96 Tissue Kit. Library preparation followed the protocol outlined in Baym et al(Baym, et al. 2015). Sequencing was carried out at Admera Health (New Jersey USA) using an Illumina HiSeq machine. Clones were sequenced using paired end 150 bp reads. In order to call mutations, the reads obtained for each clone were aligned to the *E. coli* K12 MG1655 reference genome (accession NC_000913). Mutations were recorded if they appear within a clone’s genome, but not within the ancestral genome. Alignment and mutation calling were carried out using the Breseq platform, which allows for the identification of point mutations, short insertions and deletions, larger deletions, and the creation of new junctions(Deatherage and Barrick 2014).

## Data availability statement

Metabolomics data have been deposited to the EMBL-EBI MetaboLights database, under the identifier MTBLS3041.

Raw sequencing reads were deposited to the sequence read archive (SRA) BioProject ID: PRJNA746737.

## Acknowledgements

This work was supported by ISF grants (No. 1860/21, to R.H and No. <u>824/19</u>, to E.G) and by the Rappaport Family Institute for Research in the Medical Sciences (to R.H). C.GI was supported by a TICC -Rubenstein Postdoctoral Fellowship.

---

**Figure S1.**
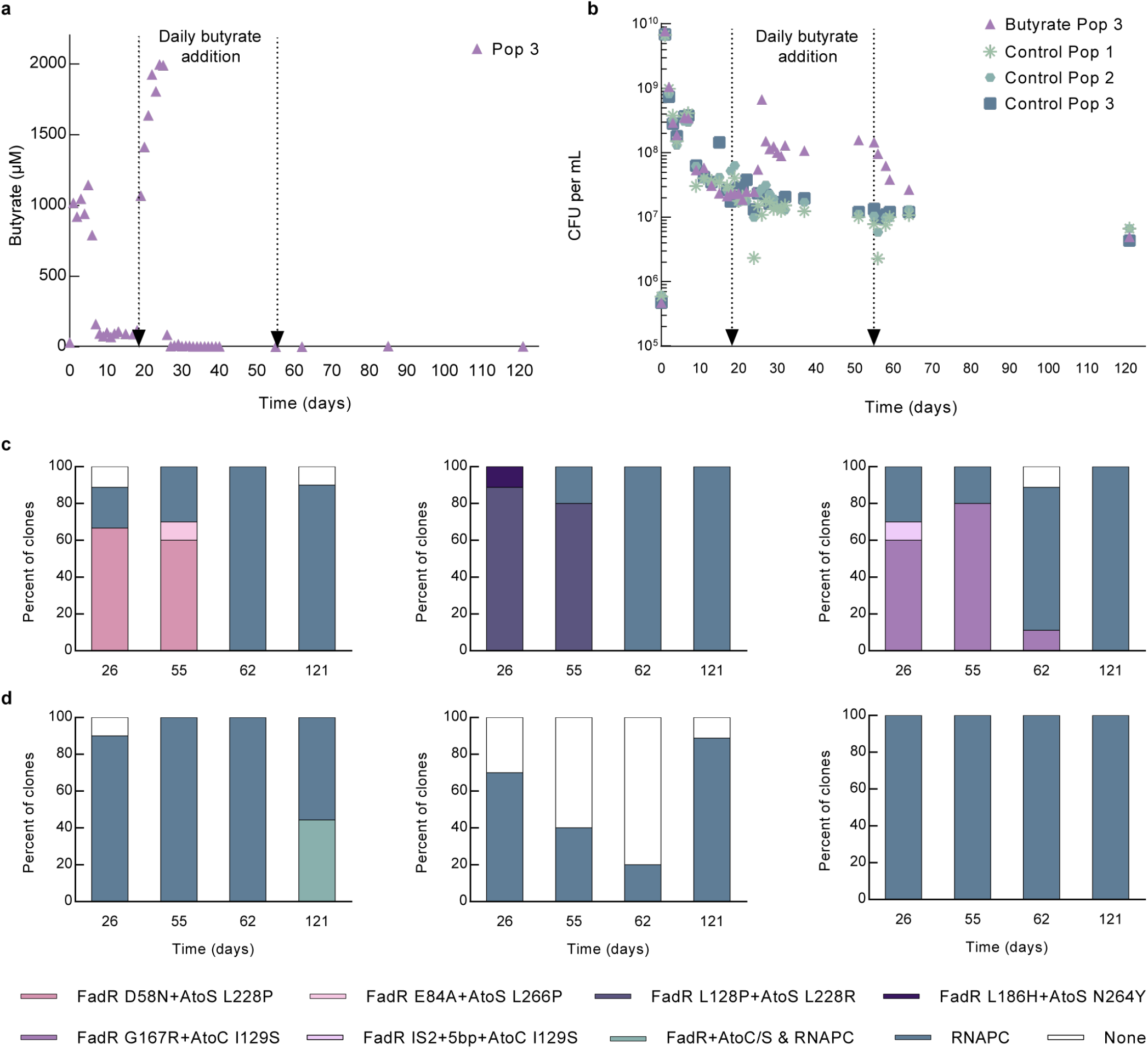
Daily butyrate supplementation and its effect on LTSP populations. **(A)** Levels of butyrate and **(B)** Mean number of viable cells, as measured through CFU calculations for the third butyrate-supplemented population. As can be seen, as with the remaining two populations (Figure 3A and 3B), for this population as well, CFU increases, once butyrate begins to be consumed. Dashed lines mark the time frame (days 18-55) during which butyrate was daily added to the media. **(C)** The frequency of different genotypes within the three populations to which butyrate was added between days 18 and 55. **(D)** The frequency of different genotypes within the three control populations to which butyrate was not added.

**Figure S2.**
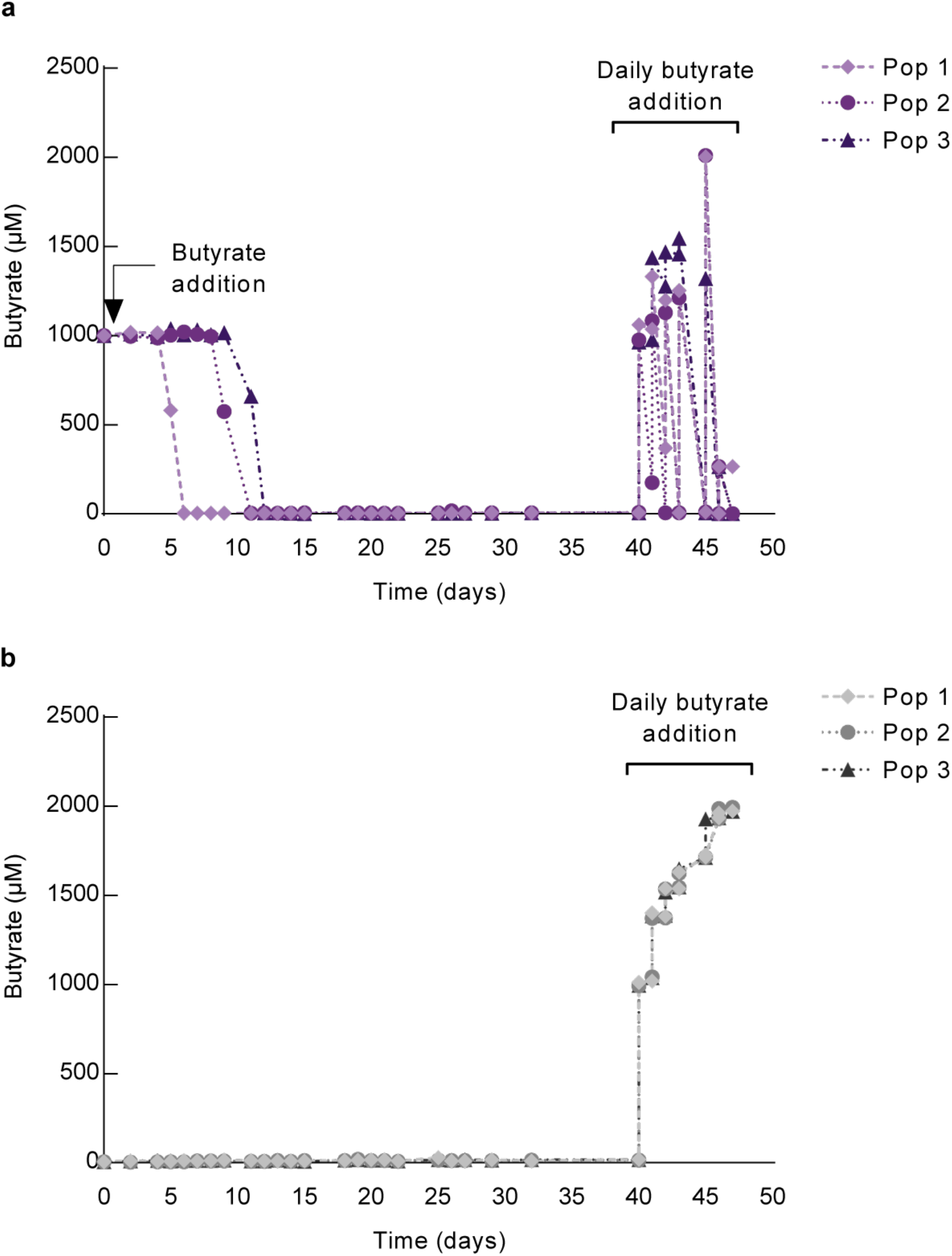
Populations that previously encountered butyrate adapt more rapidly to consume it, once they again encounter butyrate. One day after initiation, cells from three LTSP experiments were filtered out of their LB media and re-inoculated into three flasks of M9 minimal media, either supplemented with 1 mM butyrate **(A)**, or not **(B)**. When butyrate was provided, it was initially consumed by day 12. Following four additional weeks, populations were sampled and assayed for their ability to consume butyrate, which was supplemented daily into the sampled populations. As shown, only populations that initially received butyrate **(A)** could then consume it, during the week following its addition into their media (arrows represent time points in which butyrate was added).

