## Supplementary material for "Population level genetic memory of prior metabolic adaptation in *E. coli*": Figure S1

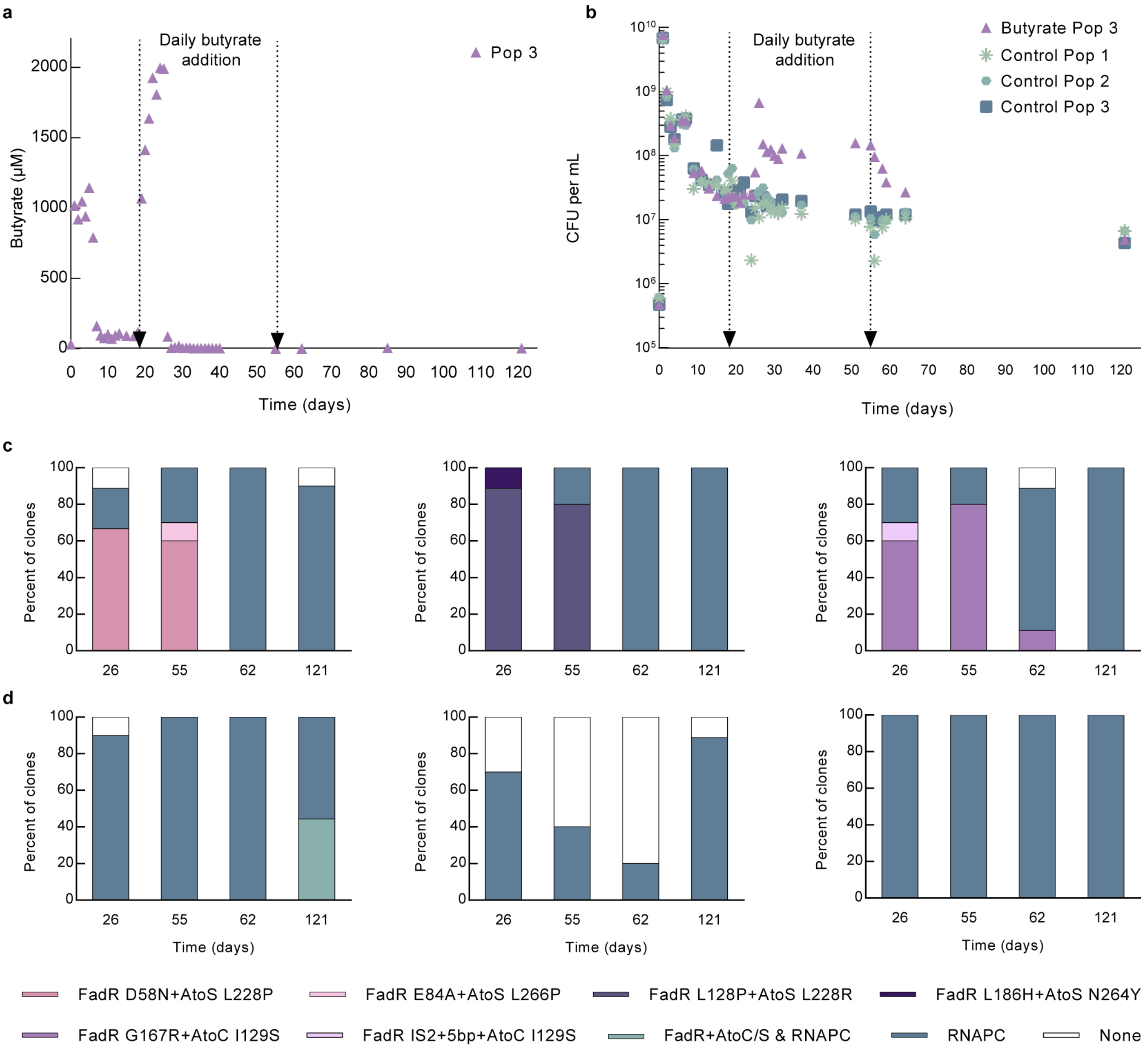


**Figure S1. Daily butyrate supplementation and its effect on LTSP populations.**

**(A)** Levels of butyrate and **(B)** Mean number of viable cells, as measured through CFU calculations for the third butyrate-supplemented population. As can be seen, as with the remaining two populations (Figure 3A and 3B), for this population as well, CFU increases, once butyrate begins to be consumed. Dashed lines mark the time frame (days 18-55) during which butyrate was daily added to the media. **(C)** The frequency of different genotypes within the three populations to which butyrate was added between days 18 and 55. **(D)** The frequency of different genotypes within the three control populations to which butyrate was not added.
