## Supplementary material for "Population level genetic memory of prior metabolic adaptation in *E. coli*": Figure S2

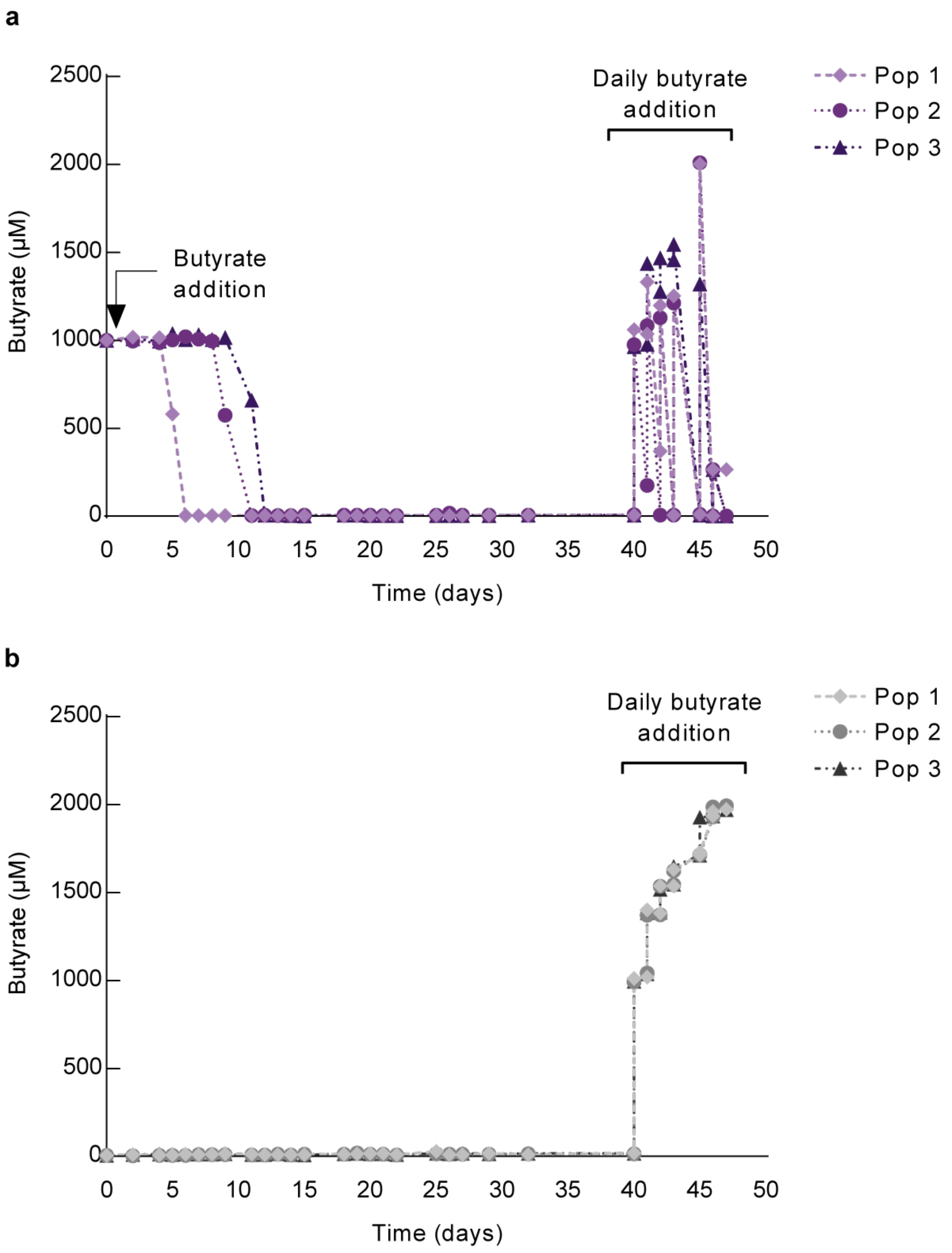


**Figure S2. Populations that previously encountered butyrate adapt more rapidly to consume it, once they again encounter butyrate**. One day after initiation, cells from three LTSP experiments were filtered out of their LB media and re-inoculated into three flasks of M9 minimal media, either supplemented with 1 mM butyrate **(A)**, or not **(B)**. When butyrate was provided, it was initially consumed by day 12. Following four additional weeks, populations were sampled and assayed for their ability to consume butyrate, which was supplemented daily into the sampled populations. As shown, only populations that initially received butyrate **(A)** could then consume it, during the week following its addition into their media (arrows represent time points in which butyrate was added).
